# Single-nucleus transcriptomic atlas of spinal cord neuron in human

**DOI:** 10.1101/2021.09.28.462103

**Authors:** Donghang Zhang, Yiyong Wei, Jin Liu, Hongjun Chen, Jin Li, Tao Zhu, Cheng Zhou

## Abstract

Despite the recognized importance of spinal cord in sensory processing, motor behaviors and/or neural diseases, the underlying neuronal clusters remain elusive. Recently, several studies attempted to define the neuronal types and functional heterogeneity in spinal cord using single cell and/or single-nucleus RNA-sequencing in varied animal models. However, the molecular evidence of neuronal heterogeneity in human spinal cord has not been established yet. Here we sought to classify spinal cord neurons from human donors by high-throughput single-nucleus RNA-sequencing. The functional heterogeneity of identified cell types and signaling pathways that connecting neuronal subtypes were explored. Moreover, we also compared human results with previous single-cell transcriptomic profiles of mouse spinal cord. As a result, we generated the first comprehensive atlas of human spinal cord neurons and defined 18 neuronal clusters. In addition to identification of the new and functionally-distinct neuronal subtypes, our results also provide novel marker genes for previously known neuronal types. The comparation with mouse transcriptomic profiles revealed an overall similarity in the cellular composition of spinal cord between the two species. In summary, these results illustrate the complexity and diversity of neuronal types in human spinal cord and will provide an important resource for future researches to explore the molecular mechanism underlying several spinal cord physiology and diseases.

## Introduction

The spinal cord is composed of distinct cell populations (Abraira et al., 2017). The heterogeneity of the various neuronal clusters contributes to the sensory perception, transduction and processing, as well as the modulation of motor behaviors (Abraira et al., 2017; Bourane et al., 2015; Mishra and Hoon, 2013; Todd, 2010). Traditionally, neuronal types are subjectively classified based on the neuronal size or established markers that related to functional aspects (Ju et al., 1987; Li et al., 2011), which cannot identify all neuronal types because of the absence of definitive markers. Moreover, functional heterogeneity among somatosensory neurons is not always in line with marker-based classification. Recently, single-cell and single-nucleus RNA sequencing provide an unbiased comprehensive strategy to explore gene expression at high resolution as well as to classify neuronal clusters in an objective manner (Aldinger et al., 2021; Li et al., 2016). Compared to single-cell RNA-seq, single-nucleus analysis gains several advantages, such as accurate cell-type analysis, easy performance in whole tissue and avoiding experimental artifacts in intact cells that induced during the tissue dissociation process (Grindberg et al., 2013; Habib et al., 2017; Lake et al., 2016).

Emerging studies have used single cell or nucleus RNA-seq to classify the neuronal types in spinal cord according to the transcriptional profiles. By Split Pool Ligation-based Transcriptome single-nucleus sequencing (SPLiT-seq), Rosenberg et al. identified 30 neuronal types in the developing spinal cord of mouse (Rosenberg et al., 2018). Delile et al. performed high-throughput single-cell RNA-sequencing and revealed spatial and temporal dynamics of gene expression in the cervical and thoracic spinal cord of developing mouse (Delile et al., 2019). Sathyamurthy et al. used massively parallel single nucleus RNA sequencing and identified 43 neuronal populations in the lumbar spinal cord of adult mouse (Sathyamurthy et al., 2018). Although the molecular and cellular organization of spinal cord is relatively well understood in mice, no research has defined the heterogeneity of neuronal types in human spinal cord by the transcriptional profile of individual cells, which is important for understanding the molecular basis for the spinal cord diseases and physiology because multiple findings from animal experiments cannot be directly replicated in human (Kushnarev et al., 2020; Yekkirala et al., 2017).

In the present study, by 10X Genomics single-nucleus RNA-sequencing, we systematically mapped the molecular and cellular compositions of human spinal cord, and compared these data with previously published mouse datasets. As a result, we classify human spinal cord neurons into 18 neuronal types with distinct transcriptional patterns, molecular markers and functional annotations. The functions of varied neuronal types in spinal cord were explored according to the transcriptome data. The resulting cell atlas demonstrates that the overall molecular and cellular organization of human spinal cord is similar with those reported in mice, although heterogeneity of transcriptional patterns exists in several clusters. The detailed information of specific gene expression may serve as an important resource of the cellular and molecular basis for physiology and etiology of spinal cord, such as disorders associated with spinal somatic sensation and motor behaviors.

## Results

### Identification of cell types

We performed single-nucleus RNA-sequencing on total 15,811 nuclei from the lumber spinal cord of two adult human (both males, 38 and 46 years old, respectively) (Figure 1A). As a result, 7 major cell types (Figure 1B) were identified with distinct molecular markers (Figure 1C-D, Supplementary Figure 1A-B): oligodendrocytes (50.6% of total nuclei), microglia (14.7% of total nuclei), neurons (13.7% of total nuclei), astrocytes (11.1% of total nuclei), oligodendrocyte precursor cells (7.8% of total nuclei), vascular cells (1.6% of total nuclei), and Schwann cells (0.5% of total nuclei). The number of genes detected per nucleus varied among these major cell types (Supplementary Figure 1C). Averagely, 5593 genes and 19398 unique molecular identifiers (UMIs) can be detected in a single DRG neuron and 2143 genes and 4355 UMIs in a single non-neuron cell.

**Figure 1.**
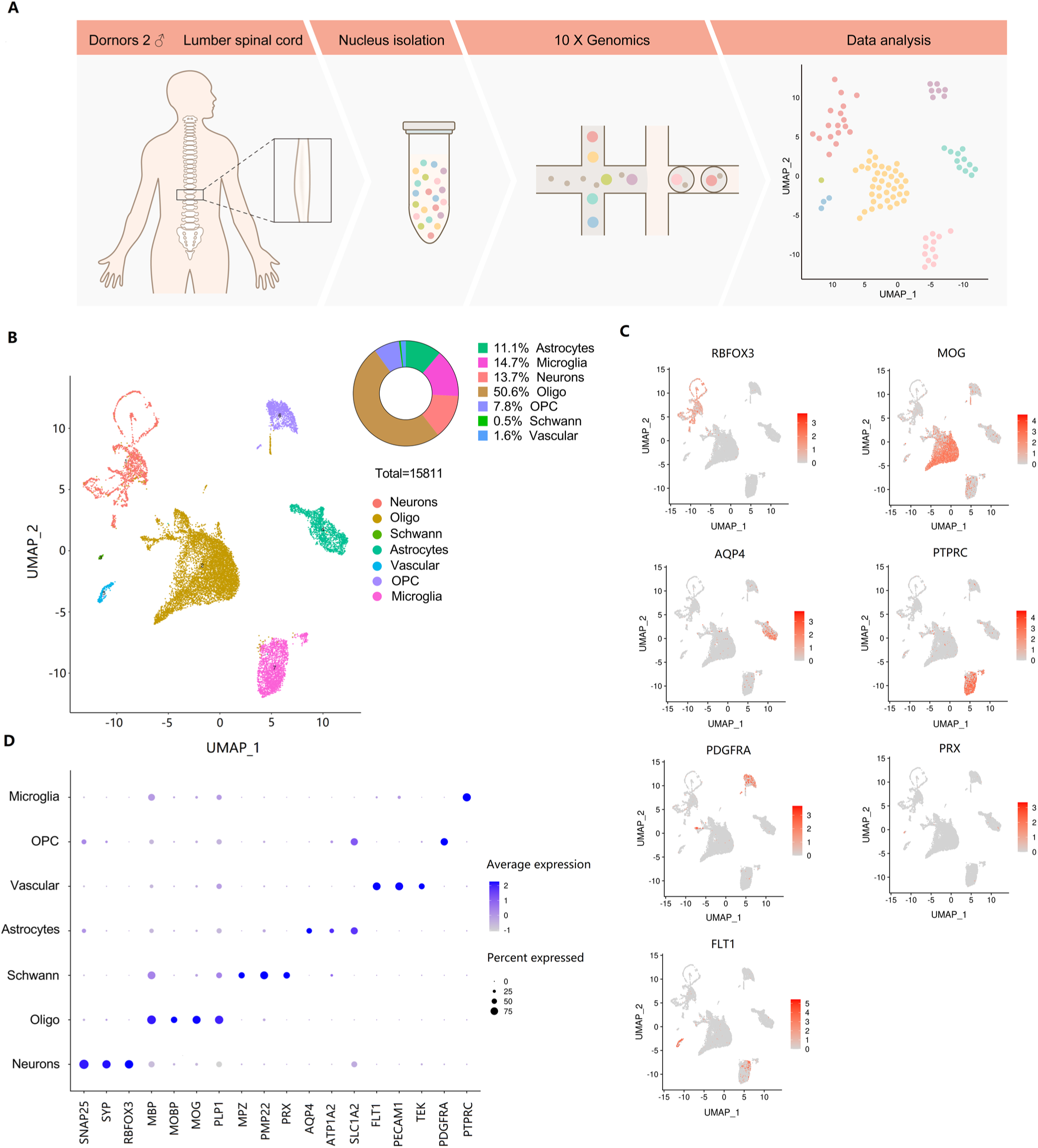
Identification of cell types. (A) Overview of the experimental workflow. (B) UMAP plot of 15,811 spinal cord cells showing 7 major cell types. Dots, individual cells; colors, cell types. (C) UMAP plot of 7 major cell populations showing the expression of representative well-known marker genes. Numbers reflect the number of UMI detected for the specified gene for each cell. (D) Dot plot showing the distribution of expression levels of selected marker genes across all 7 cell types.

### Identification of neuronal subtypes

To identify and characterize neuronal subtypes within the spinal cord, 2163 neuronal nuclei were unbiasedly reclassified into 18 subclusters (H1-H18) (Figure 2A) based on their transcriptional characteristics. These neuronal clusters were further grouped into 2 different functional clusters: 10 clusters of excitatory neurons (glutamatergic, with markers of SLC17A6), and 8 clusters of inhibitory neurons (GABAergic/glycinergic, with markers of GAD1, GAD2 and SLC6A5) (Figure 2B) with distinct neurotransmitter markers (Figure 2C-D, Supplementary Figure 2A-B). In total, 50.7% of neuronal nuclei were in predominantly excitatory clusters (including 6.3% motor neurons with cholinergic markers of CHAT and SLC5A7), and 49.3% were in predominantly inhibitory clusters.

**Figure 2.**
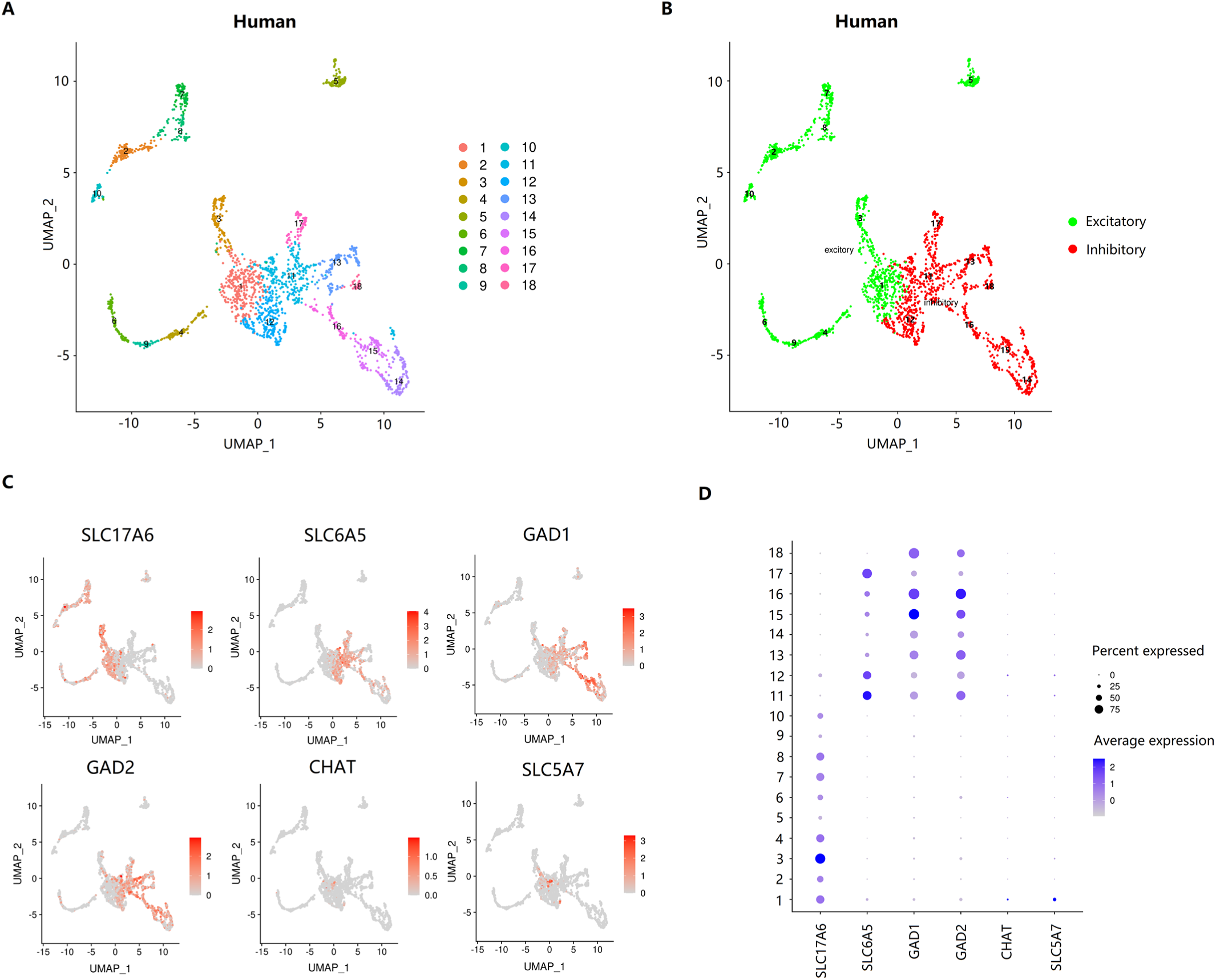
Identification of neuronal subtypes in human spinal cord. (A) UMAP plot of 2163 spinal cord neurons showing 18 neuronal clusters. Dots, individual cells; colors, neuronal clusters. (B) UMAP plot of spinal cord neurons based on the excitatory and inhibitory marker genes. Dots, individual cells; colors, neuronal clusters. (C) UMAP plot of excitatory and inhibitory clusters showing the expression of representative well-known marker genes. Numbers reflect the number of UMI detected for the specified gene for each cell. (D) Dot plot showing the distribution of expressional levels of selected marker genes across all the 18 neuronal clusters.

### Human-mouse cell type homology

To determine the similarities and differences of neuronal subtypes between human and mouse spinal cord, we reclassified the mouse spinal cord neurons into 23 clusters (M1-M23) (Figure 3A) (including 13 excitatory clusters and 10 inhibitory clusters) (Figure 3B-D, Supplementary Figure 2C-D) using published datasets by single nuclei sequencing from the mouse spinal cord (Sathyamurthy et al., 2018). As a result, 44.7% of neuronal nuclei (total 4280) were distributed in excitatory clusters (including 3.5% motor neurons) and 55.3% were in inhibitory clusters.

**Figure 3.**
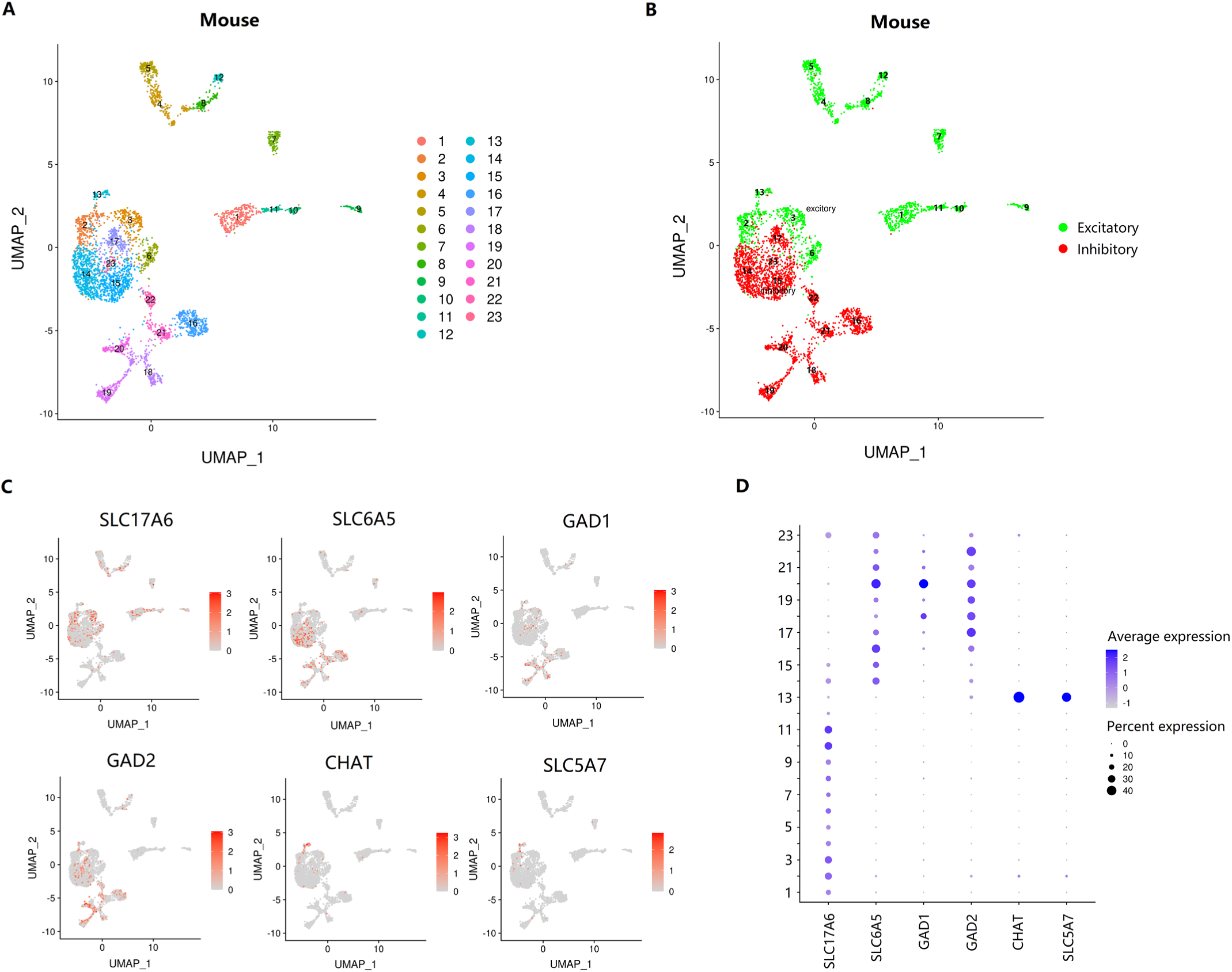
Identification of neuronal subtypes in mouse spinal cord. (A) UMAP plot of 4280 spinal cord neurons showing 23 neuronal clusters. Dots, individual cells; colors, neuronal clusters. (B) UMAP plot of spinal cord neurons based on the excitatory and inhibitory marker genes. Dots, individual cells; colors, neuronal clusters. (C) UMAP plot of excitatory and inhibitory clusters showing the expression of representative well-known marker genes. Numbers reflect the number of UMI detected for the specified gene for each cell. (D) Dot plot showing the distribution of expressional levels of selected marker genes across all the 23 neuronal clusters.

Next, we used co-clustering methods (Stuart et al., 2019) to align the identified cell types in human spinal cord with that from the mouse spinal cord. Overall, the neuronal clusters of human spinal cord grouped well with corresponding mouse counterparts in the UMAP (Figure 4A-B). Although UMAP plots provide a visual representation of similarity between cells with related transcriptomic properties, relationships between separated clusters are hard to interpret due to the collapsed multidimensional information into two dimensions. Therefore, we used Kullback-Leibler divergence (KLD) (Perez-Cruz, 2008) estimation to quantitate the similarity between human neuronal clusters and their potential mice counterparts. The quantitative results revealed that, the closed clusters of human and mice in the UMAP showed strong concordance as compared to other clusters (Figure 4C). Finally, we defined the homology clusters between human and mice according to the alignment results, including H1-M2, M14, H2-M8, H3-M2, M3, M6, M13, H4-M1, H5-M7, H6-M9,10, H7-M5, H8-M4, H9-M11, H10-M12, H11-M14, M15, M17, M23, H12-M14, H13-M16, M21, H14-M19, H15-M18, H16-M20, H17-M22, H18-M18 (Figure 4D).

**Figure 4.**
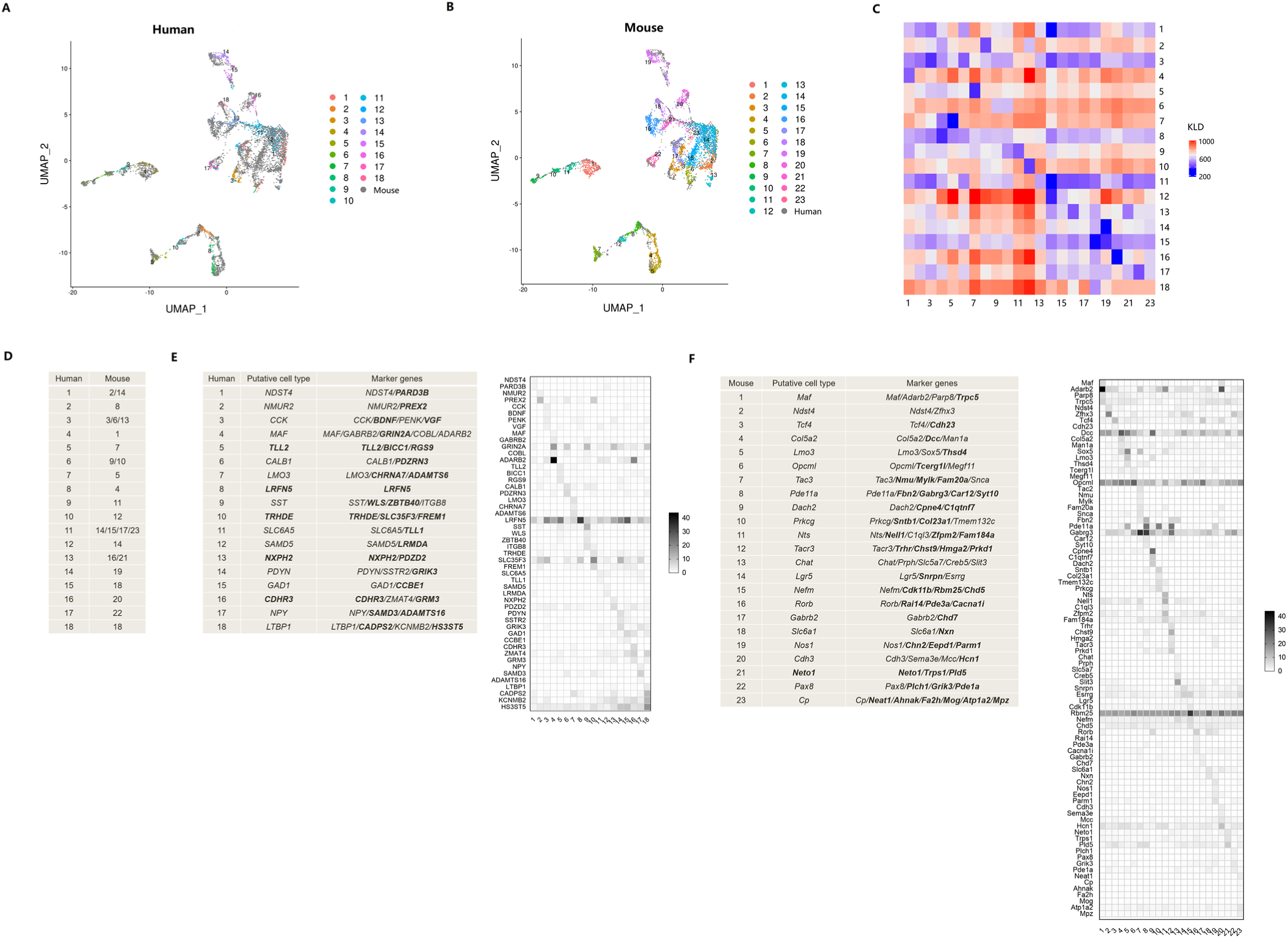
Human-mouse cell type homology. (A) UMAP plot of the co-clustering of mouse and human neurons. Dots, individual cells; colors, human neuronal clusters. (B) UMAP plot of the co-clustering of mouse and human neurons. Dots, individual cells; colors, mouse neuronal clusters. (C) Heatmap showing the natural logarithm of Kullback-Leibler divergences (KLD) for human neuronal clusters versus mouse neuronal clusters. The similarity is inversely proportional to the KLD values. (D) The putative homologous neuronal clusters between human and mouse. (E-F) Putative neuronal type and their representative marker genes in human (E) and mouse (F). Previously undescribed neuron types and marker genes are shown in bold. Heatmap showing the expressional levels of the marker genes across different clusters.

### Predominant markers in neuronal clusters

After classification of neuronal clusters, we sought to explore the representative marker genes and then annotate these clusters. To be considered as a marker gene, the prerequisites are selectivity, a relatively high level of expression, and utility in multiple experimental approaches. Previously known markers can define most of these clusters, while we also provide new marker genes (Figure 4E-F). Additionally, we identified several previously unrecognized neuronal populations (Figure 4E-F).

To examine the conservation of molecular and cellular architecture in the spinal cord between human and mouse, we compared the gene profiles between human neuronal clusters and all their putative mouse counterparts. We found that shared genes define several neuronal clusters in both species, such as H1-M2, H4-M1, H7-M5, H16-20 (Figure 4E-F). Furthermore, the homologous clusters between human and mouse both show high expression values of several same genes (Supplementary Figure 3A-B). For example, the homologous clusters H4 and M1 both show high expression values for ADARB2/Adarb2 (adenosine deaminase RNA specific B2), ERBB4/Erbb4 (erb-b2 receptor tyrosine kinase 4), ARPP21/Arpp21 (cAMP regulated phosphoprotein 21), and GRIN2A/Grin2A (glutamate ionotropic receptor NMDA type subunit 2A).

Then, we compared the expression profiles of classical spinal cord markers between the homologous clusters of human and mouse, including ion channels, neurotransmitter receptors, neuropeptides, and transcription factors (Figure 5). We firstly compared the expression levels of genes between different clusters of human and/or mouse. The results showed that several genes are differentially expressed among different neuronal clusters. For example, the expression of SCN7A (Na_v_2.1) is lower in cluster H2 compared to other clusters of humans (Figure 5A). The expression of HCN1 (hyperpolarization activated cyclic nucleotide gated potassium channel 1) is higher in cluster M20 compared to other clusters of humans (Figure 5D). The expression of GRIN2A (glutamate ionotropic receptor NMDA type subunit 2A) is lower in clusters H2 and H7 compared to other clusters of humans (Figure 6A). Compared to other clusters of humans, the expression of CHRNA7 (cholinergic receptor nicotinic alpha 7 subunit) (Figure 6A) and NPY (neuropeptide Y) (Figure 6C) are extremely higher in cluster H7 and H17, respectively.

**Figure 5.**
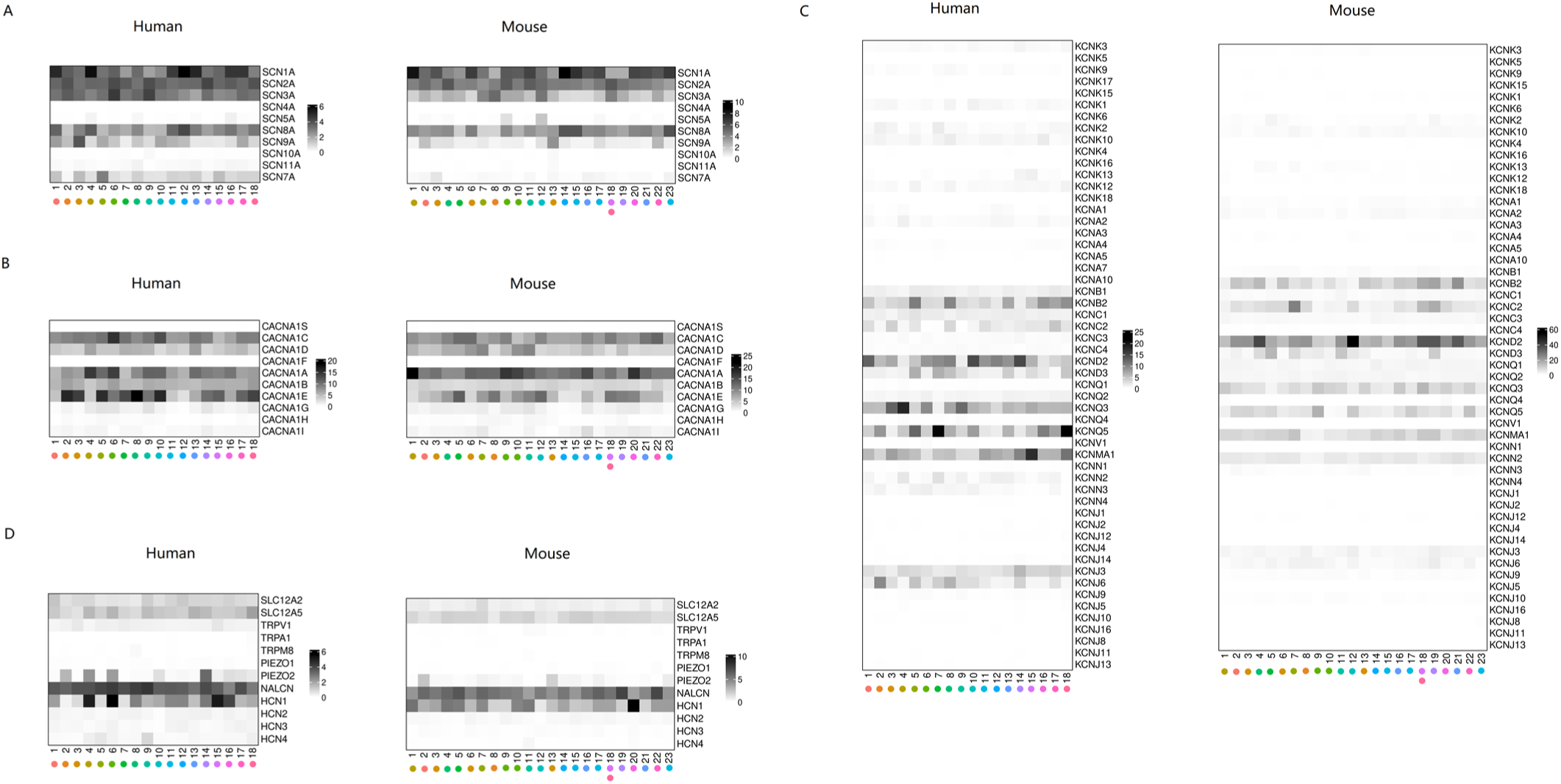
The expressional profiles of classical marker genes for ion channels in human and mouse. (A-D) Normalized mean gene expression of classic marker genes of sodium channels (A), calcium channels (B), potassium channels (C), and other ion channels (D). The same color showing the homologous clusters of humans and mice.

**Figure 6.**
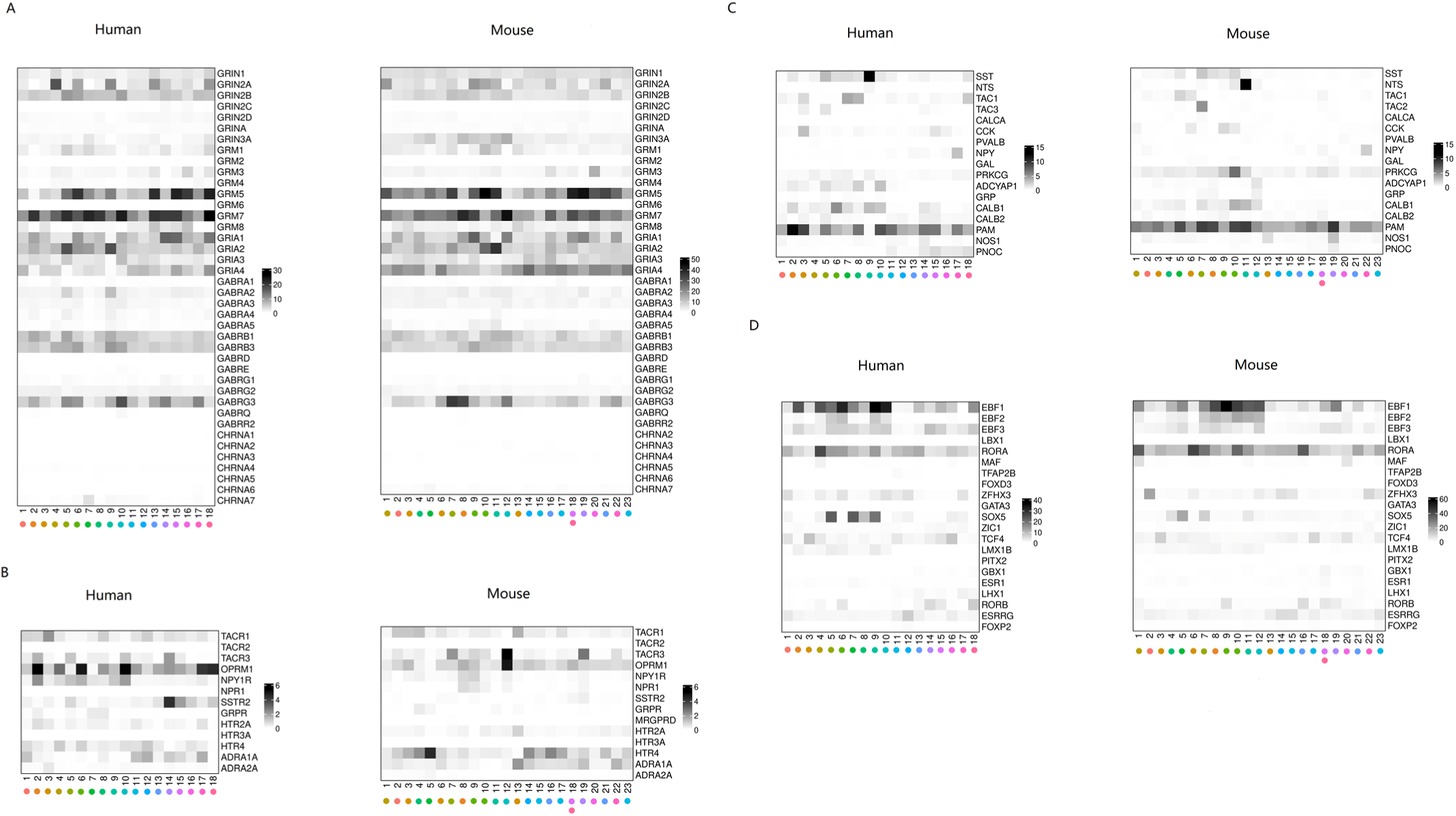
The expressional profiles of classical marker genes for receptors, neuropeptides, and transcription factors in human and mouse spinal cord. (A-D) Normalized mean gene expression of classic marker genes of glutamatergic-, GABAergic-, and cholinergic receptor (A), other receptors (B), neuropeptides (C), and transcription factors (D). The same color showing the homologous clusters of humans and mice.

Next, we compared transcriptional patterns of classical marker genes in the putative homologous clusters between human and mouse. The overall transcriptional patterns are similar between human and mouse spinal cord. For example, SCN5A/Scn5a (sodium voltage-gated channel alpha subunit 5, Na_v_1.5) is present in the M12 cluster but is very low-expression in the homologous H10 cluster (Figure 5A). KCNK1/Kcnk1 (potassium two pore domain channel subfamily K member 1, TWIK1) is present in multiple clusters of humans but is very low-expression in all clusters of mice (Figure 5C). TRPV1/Trpv1 (transient receptor potential cation channel subfamily V member 1) is widely present in the human spinal cord but is relatively low-expression in all clusters of mice (Figure 5D).

For classical receptors, SSTR2/Sstr2 (somatostatin receptor 2) (Figure 6B) are widely present in the neuronal clusters of human spinal cord but is very low-expression in most clusters of mice. NPR1/Npr1 (natriuretic peptide receptor 1) is present in several neuronal clusters of mouse spinal cord but very low-expression in all clusters of humans (Figure 6B). HTR4/Htr4 (5-hydroxytryptamine receptor 4) is present in the M19 cluster but not in the putative homologous cluster of H14 (Figure 6B).

For neuropeptides, SST/Sst (somatostatin) shows high expression levels in H9 but not in the homologous cluster of M11 (Figure 6C). PNOC/Pnoc (prepronociceptin) is widely present in the neuronal clusters of H14-18 but is very low-expression in almost all clusters of mice (Figure 6C). For transcription factors, EBF2/Ebf2 (EBF transcription factor 2) is present in M4 and M5 but not in their homologous clusters of H8 and H7, respectively (Figure 6D).

### Functional assignment for neuronal clusters

To propose a relation of the identified neuronal types to known modality-specific functions, we performed Gene ontology (GO) term analysis of their top genes. We found that the overall function assignment for excitatory clusters was similar with that for inhibitory clusters in human (Supplementary Figure 4A-B) or mouse (Supplementary Figure 4C-D). And the function assignment for excitatory and inhibitory clusters was similar between human and mouse (Supplementary Figure 4A-D). In detail, synaptic transmission, neurotransmitter receptors activities of glutamate, GABA and NMDA, ion transport of potassium, sodium and calcium, axon guidance and neuron projection development components are significantly over-represented among the top genes that contribute to defining human excitatory clusters (Supplementary Figure 4A); while synaptic transmission, glutamate receptor- and chemokine-mediated signaling pathway, axon guidance and axonogenesis, central nervous development, calcium ion transport, glial cell differentiation components are significantly over-represented among the top genes that contribute to defining human inhibitory clusters (Supplementary Figure 4B). In the mouse, glutamate receptor- and netrin-activated signaling pathway, calcium and sodium ion transport, locomotor behavior and behavioral fear response, synaptic transmission, neuron migration, axon guidance, dendrite morphogenesis and nervous development components are significantly over-represented among the top genes that contribute to defining excitatory clusters (Supplementary Figure 4C); while glutamate receptor- and acetylcholine signaling pathway, synaptic transmission, cell adhesion, axon guidance and neuron morphogenesis, calcium ion transport, locomotor behavior, and neuromuscular junction development components are significantly over-represented among the top genes that contribute to defining inhibitory clusters (Supplementary Figure 4D).

### Cellular convergence of disease

Neuronal dysfunction within spinal cord underlies multiple complex disorders. As a framework for understanding these complex disorders in human, we used our atlas of spinal cord to identify the neuron types in which changes of gene expression may be associated with these diseases. We first examined the transcriptional profiles of genes implicated in two common types of chronic pain, namely neuropathic pain (Figure 7A) and inflammatory pain (Figure 7B). Several risk genes of neuropathic pain in human, including GRM5 (glutamate metabotropic receptor 5, G-protein coupled receptor), GRIN2B (glutamate ionotropic receptor NMDA type subunit 2B, Ion channel), OPRM1 (opioid receptor mu 1, G-protein coupled receptor), HMGB1 (high mobility group box 1, Nucleic acid binding), MCF2L2 (MCF.2 cell line derived transforming sequence-like 2, Signaling), and DNMT3A (DNA methyltransferase 3 alpha) show a higher expression level in most neuronal clusters as compared to the rest genes (Figure 7A left). Cluster H7 shows predominant expression of BDNF (brain derived neurotrophic factor, Signaling) and SCN9A (sodium voltage-gated channel alpha subunit 9, Na_v_1.7) (Figure 7A left). TAC1 (tachykinin precursor 1) is present at high levels in clusters H7, H12, H15, and H18 compared to other clusters of humans (Figure 7A left). Then we examined the expression of these risk genes in mouse spinal cord. We found that the homologous genes Grm5, Grin2b, and Dnmt3a also have a higher expression level in most clusters (Figure 7A right). Oprm1 shows higher expression in clusters H21, and Scn9a shows higher expression in cluster H5, H17, and H22 (Figure 7A right).

**Figure 7.**
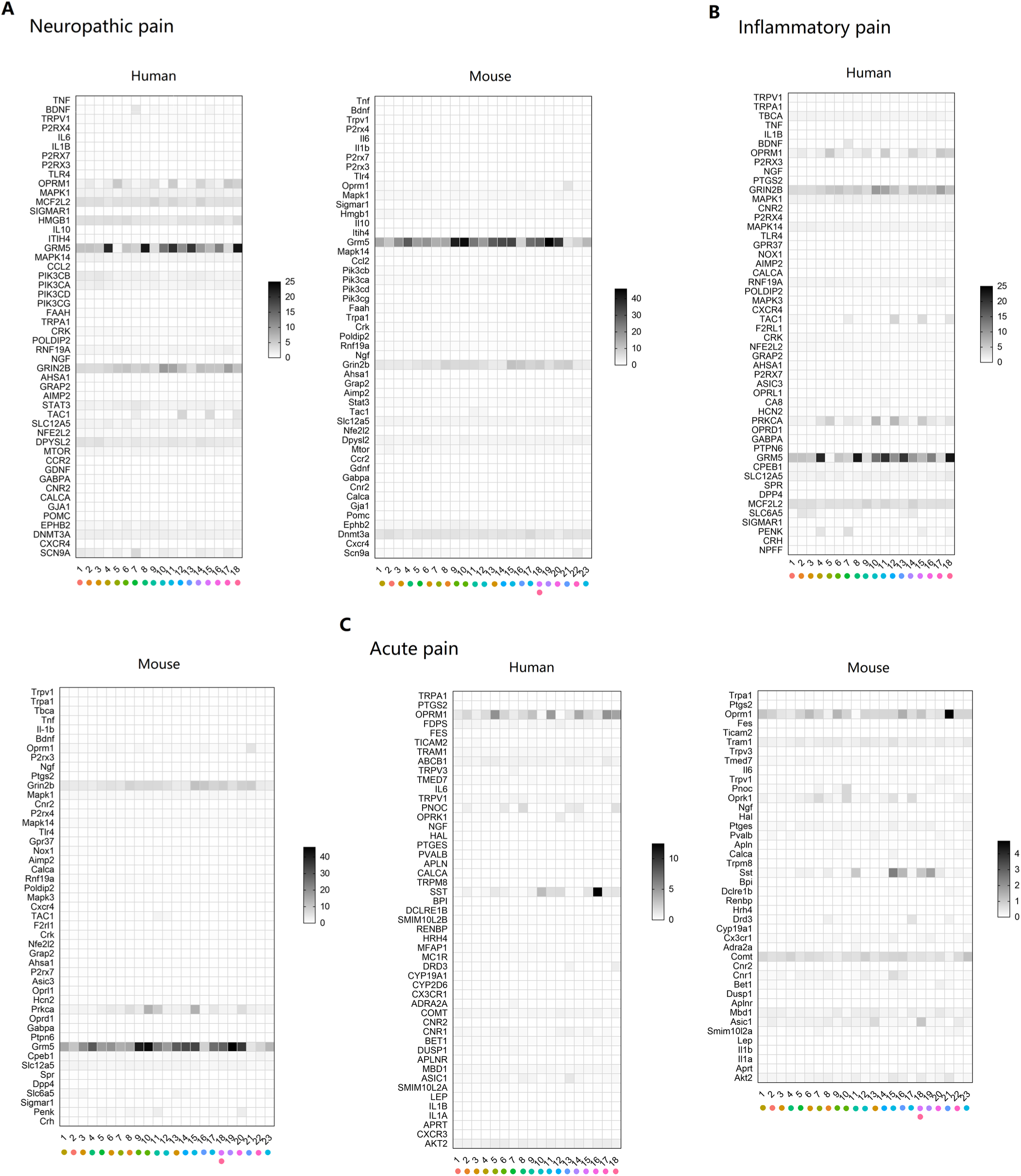
The expressional profiles of risk genes for spinal cord diseases in human and mouse. (A-C) The transcriptional profiles of risk genes for neuropathic pain (A), inflammatory pain (B), and acute pain (C). The same color showing the homologous clusters of humans and mice.

The predominant risk genes of neuropathic pain also contribute to inflammatory pain in human. GRM5, GRIN2B, MCF2L2, OPRM1, and PRKCA (protein kinase C alpha, Kinase) shows a higher expression level in most clusters than other genes (Figure 7B left). BDNF shows higher expression in cluster H7 (Figure 7B left). TAC1 shows higher expression in clusters H7, H12, H15, and H18 (Figure 7B left). SLC6A5 (solute carrier family 6 member 5, Transporter) shows higher expression in clusters H2, H3, H13, and H14 (Figure 7B left). PENK (proenkephalin, Signaling) shows higher expression in clusters H4 and H7 (Figure 7B left). In mouse spinal cord, the homologous genes Grm5, Grin2b, Prkca shows a higher expression level in most clusters (Figure 7B right). Oprm1 shows higher expression in cluster H21 (Figure 7B right). Penk shows higher expression in clusters H11 and H20 (Figure 7B right).

Then, we examined the expression of genes that cause acute pain in human (Figure 7C). OPRM1, SST (somatostatin, Signaling), AKT2 (AKT serine/threonine kinase 2, Kinase), and COMT (catechol-O-methyltransferase, Enzyme) exhibits higher expression level in most clusters (Figure 7C left). PNOC (prepronociceptin, Signaling) shows higher expression in clusters 6, 8, H13, H14 and H18 (Figure 7C left). OPRK1 (opioid receptor kappa 1, G-protein coupled receptor) shows higher expression in cluster H12 (Figure 7C left). DRD3 (dopamine receptor D3, G-protein coupled receptor) shows higher expression in cluster H18 (Figure 7C left). ASIC1 (acid sensing ion channel subunit 1, Ion channel) shows higher expression in cluster H13 (Figure 7C left). The homologous genes Oprm1, Sst, Comt, and Oprk1 show a higher expression level in most clusters (Figure 7C right). Pnoc shows higher expression in cluster H10 (Figure 7C right). Drd3 expresses higher in cluster H17 and Asic1 shows higher expression in cluster H13 and H18 (Figure 7C right).

Next, we showed the transcriptional profiles of risk genes of other spinal cord disease in human and mouse spinal cord, including 3 types of inflammatory diseases (Multiple sclerosis, Herpes zoster, and Poliomyelitis) (Figure 7D, Supplementary Figure 5), 3 types of developmental abnormalities (Diastematomyelia, Syringomyelia, and Hereditary ataxias) (Supplementary Figure 5), and one type of motor neuron disease, namely Amyotrophic lateral sclerosis (ALS) (Figure 7E).

Taken together, these findings demonstrate the value of our spinal cord atlas as a rich resource for probing the cellular biology underlying complex disease in spinal cord.

## Discussion

A large number of evidence from animal’s studies demonstrate the diversity and complexity of cells composition in spinal cord (Rosenberg et al., 2018; Sathyamurthy et al., 2018). In spinal cord, the heterogeneity of the various neuronal components makes up neuronal circuits that process sensory information and regulate motor behavior (Abraira et al., 2017; Bourane et al., 2015; Todd, 2010). To our knowledge, there is no study has shed light on the neuronal organization of spinal cord in human. In the present study, we firstly profiled the classification of cell types in human spinal cord by 10X Genomics single-nucleus RNA sequencing. As a result, we created an atlas of 18 discrete neuron subtypes in human spinal cord. We also defined these neuronal clusters with representative marker genes.

To date, multiple cellular and molecular changes in spinal cord, including ion channels, neurotransmitter receptors, neuropeptides, and transcription factors, are thought to contribute to pain development after nerve injury (Yekkirala et al., 2017). Hence, the spinal cord provides numerous potential targets for the development of novel analgesics. However, it is a great challenge to transfer the experimental results from animals directly to humans (Kushnarev et al., 2020; Tibbs et al., 2016; Yekkirala et al., 2017). High variance in cell components and gene expression of individual cell type in spinal cord among species may partly explain the failure of clinical translation. Our results from human-mouse comparisons indicate substantial similarities in cell components and the expression profiles of important genes across species. However, several neuronal clusters showed varied expression profiles in terms to specific gene. Notably, we also identified some pain-related genes that expressed in the mouse spinal cord was not present in human. This may partially explain why drug targets in the mouse model cannot be replicated in human. Therefore, the transcriptional profiles of spinal cord neurons in other species closer to human are needed, such as non-human primate.

The spinal cord has a pivotal role in normal sensory processing and in pathological conditions, such as chronic pain, providing numerous potential targets for the development of novel analgesics (Todd, 2010). Although multiple risk genes have been identified by previous studies, the expression profiles of these risk genes in neuronal clusters of spinal cord remains elusive in human. In this study, we have provided the transcriptional patterns of risk genes of 3 types of pain, including neuropathic pain, inflammatory pain, and acute pain. We have identified the expression level of each risk gene in different neuronal clusters. We also compared the difference in the expression of risk genes across species. In addition, we checked the transcriptional patterns of risk genes of other common human diseases implicated in spinal cord, including 3 types inflammatory diseases (Multiple sclerosis, Herpes zoster, and Poliomyelitis), 3 types developmental abnormalities (Diastematomyelia, and Syringomyelia, and hereditary ataxias), and one motor neuron disease (Amyotrophic lateral sclerosis). Therefore, this work reveals the molecular repertoire of each neuronal population, providing a significant extension of our understanding of spinal cord cell types, providing an important database which allow researchers to probe and analyze the expression profiles of risk genes for human diseases revolving spinal cord. Therefore, this data resource may serve as a powerful tool to advance our understanding of the molecular mechanisms in neuronal populations that mediate spinal cord diseases.

According to the projection sites of axon, neurons in spinal cord can be divided into two main types, including projection cells with their axons that project to the brain and interneurons with their axons that remain within the section of spinal cord (Todd, 2017). The latter can be further divided into two functional classes: inhibitory neurons, which use GABA and/or glycine as their principal transmitter, and excitatory neurons, which use glutamate as the principal transmitter (Todd, 2017). In this work, we also classified the spinal neurons into these two functional clusters with excitatory and inhibitory markers, respectively. We found that rare individual neuronal nuclei co-expressed excitatory and inhibitory markers, which is consistent with the data from mouse (Sathyamurthy et al., 2018). After co-clustering of the human and mouse data, we found that the excitatory and inhibitory clusters of human maps well with that from the mouse. In accord with previous study (Todd, 2017), our results also show that, some classical marker genes of spinal cord, including CCK, SST, CALB1, Nts, Tac2, and Prkcg are predominantly present in excitatory neuronal clusters, others, such as NPY and Nos1 are predominantly present in inhibitory neuronal clusters. Previous study reported that the genes of opioid peptides Penk and Pdyn are expressed by both excitatory and inhibitory interneurons of mouse (Todd, 2017), while our results showed that PENK are predominantly present in excitatory neuronal cluster 3 and PDYN are predominantly present in inhibitory neuronal cluster 14 of human. The neurokinin 1 receptor (NK1R, the receptor for substance P), also known as TACR1, is predominately expressed on projection neurons, but is also present on excitatory interneurons (Todd, 2010, 2017). TACR1 is commonly used as the marker gene of projection neurons within spinal cord in animal research. Previous study has revealed that projection neurons only make up a very small portion of spinal cord neurons (Todd, 2010). Consistently, the transcriptional data from mouse showed that only 10% nuclei expressed TACR1. Therefore, the exact ratio of projection neurons should be less than 10%. Notably, the results here indicate that TACR1 is present in not only the excitatory clusters, but also the inhibitory neuronal clusters (Supplementary Figure 6A). The transcriptional data from human also showed that TACR1 is present in both excitatory and inhibitory neuronal clusters (Supplementary Figure 6B). But the ratio of TACR1 positive neurons is higher (44%) than that in the mouse.

There are several limitations in this work. Firstly, we did not validate the marker genes using immunohistochemical methods due to the limited samples, as well as the function of the identified neuronal clusters. Secondly, the spatial location and the connection of each neuronal cluster were not determined, which may be solved *via* spatial transcriptomics analysis. The third limitation is that the sample is small and the data is only from the males.

## Methods

### Human spinal cord samples

This study was approved by the Ethical Committee of Affiliated Hospital of Zunyi Medical University on May 19, 2021 (Approval No. KLL-2020-273) and was registered with the Chinese Clinical Trial Registry (ChiCTR2100047511, principal investigator, Yiyong Wei) on Jun 20, 2021. Written informed consent was obtained prior to patient enrolment from family numbers of patients. All experiment protocol were performed in accordance with ethical and legal guidelines. Lumber spinal cord was acutely isolated from two male adult brain-dead human donors without previous diagnosis of acute/chronic low back or lower limb pain. Samples were immediately cleaned from the blood and connective tissue, transferred into liquid nitrogen to snap freeze, and stored at −80 °C until nuclei isolation.

### Isolation of Nuclei

Nuclei were isolated using a Nucleus Isolation Kit (catalogue no. 52009-10, SHBIO, CHINA) according to the manufacturer’s instructions. Briefly, frozen human DRG and spinal cord were thawed on ice, minced and homogenized in cold 1% BSA in LB solution. Then, the lysate was filtered through a 40-μm cell strainer and was centrifuged at 500 × *g* for 5 min at 4 °C. After removing the supernatants, the pellet was resuspended in LB. PB1, PB2 and PB3 was added in sequence, followed by a centrifugation at 3000 × *g* for 20 min at 4 °C. The pellet was then filtered through a 40-μm cell strainer, centrifuged at 500g for 5 min at 4 °C, and resuspended for two times in NB. Nuclei were sorted by staining and counted with a dual-fluorescence cell counter.

### cDNA synthesis

Nuclei suspension was loaded onto the Chromium single cell controller (10x Genomics) to generate single-nucleus gel beads in the emulsion (GEM) using single cell 3’ Library and Gel Bead Kit V3.1 (10x Genomics, 1000075) and Chromium Single Cell B Chip Kit (10x Genomics, 1000074) according to the manufacturer’s protocols. Approximately 6058 nuclei were captured for each sample. The captured nucleus was lysed and the released mRNA was barcoded through reverse transcription in individual GEM. Reverse transcription was performed to generate cDNA using a S1000TM Touch Thermal Cycler (Bio Rad) with the following parameters: 53°C, 45 min; 85°C, 5 min; hold, 4°C. The cDNA was then amplified, and the quality was assessed using an Agilent 4200 (CapitalBio Technology, Beijing).

### 10x Genomics Library Preparation and Sequencing

snRNA-seq library was established with Single Cell 3’ Library and Gel Bead Kit V3.1 according to the manufacture’s introduction. The libraries were sequenced using a Novaseq 6000 sequencing platform (Illumina) with a depth of at least 30,000 reads per nucleus with pair-end 150 bp (PE150) reading strategy (CapitalBio Technology, Beijing).

### Single-nucleus transcriptomic data analysis Seurat pipeline

Cell clustering was performed using the Seurat 3.0 (R package). Cells were filtered out if the gene number was less than 200 or ranked in the top 1%, or the mitochondrial gene ratio was more than 25%. Dimensionality reduction was performed using Principal component analysis (PCA) and visualized by UMAP.

### Enrichment Analysis

Gene ontology (GO) enrichment was conducted using R package with default settings. GO terms with adjusted p-value of less than 0.05 calculated by the hypergeometric test followed by the Benjamini-Hochberg method were defined as significant enriched term. Top 15 enriched terms were selected to be visualized.

### Cell clusters annotation

The marker genes were identified by using the ‘FindAllMarkers’ function in Seurat with settings on genes with at least 0.25 increasing logFC upregulation, comparing to the remaining cells.

### Risk genes of spinal cord diseases

The risk genes of spinal cord diseases were identified using DisGeNET, which contains one of the largest publicly available collections of genes associated with human diseases. If the number of risk genes was more than 50, we only chose the top 50 risk genes ordering by the number of PMIDs.

## Supplementary figures and legends

**Supplementary Figure 1.**
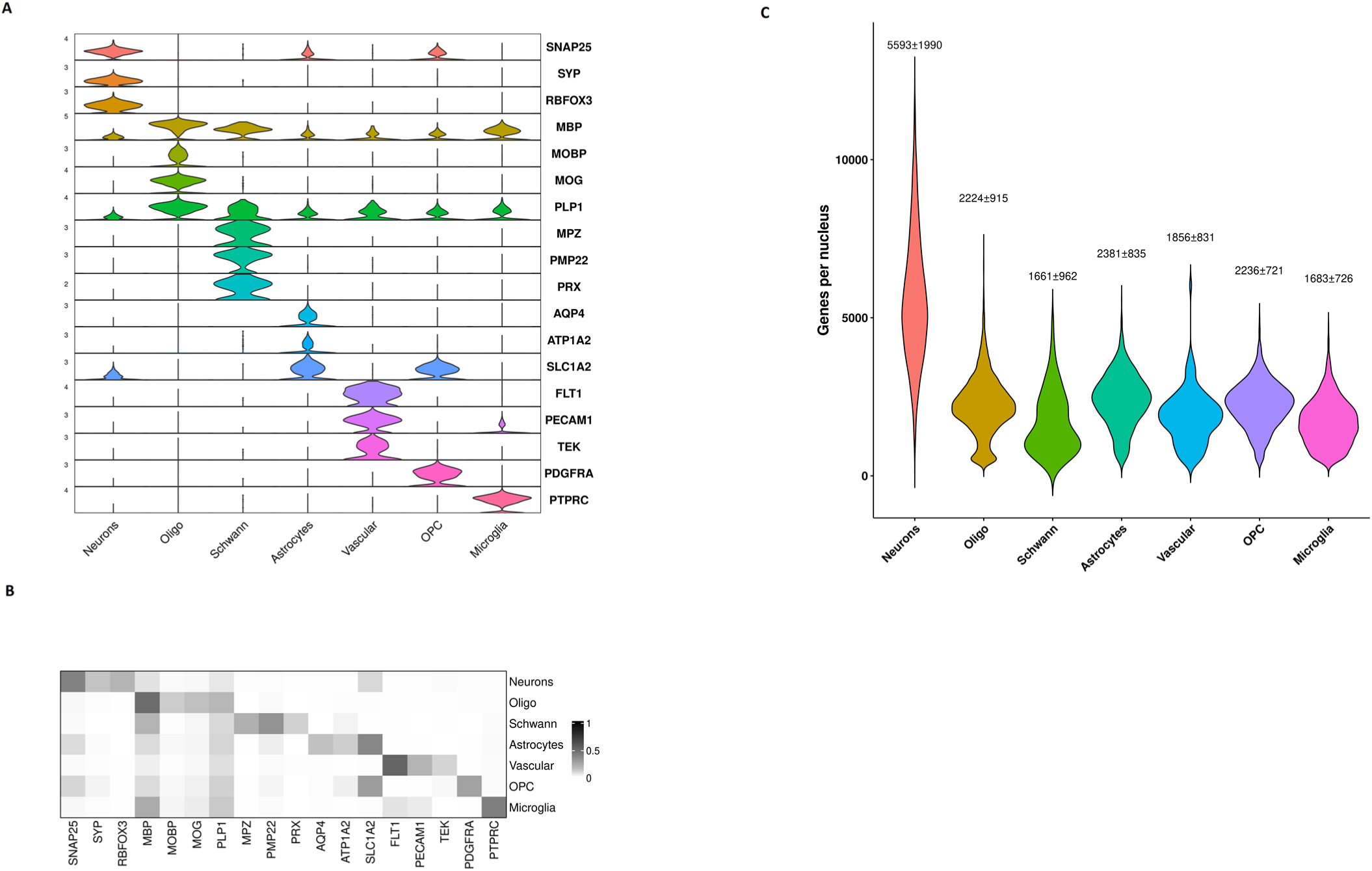
(A-B) Violin plot (A) and heatmap (B) showing the distribution of expression levels of selected marker genes across all 7 cell types. (C) Violin plot showing genes detected per nucleus in each cell type. Data is present as mean ± SD.

**Supplementary Figure 2.**
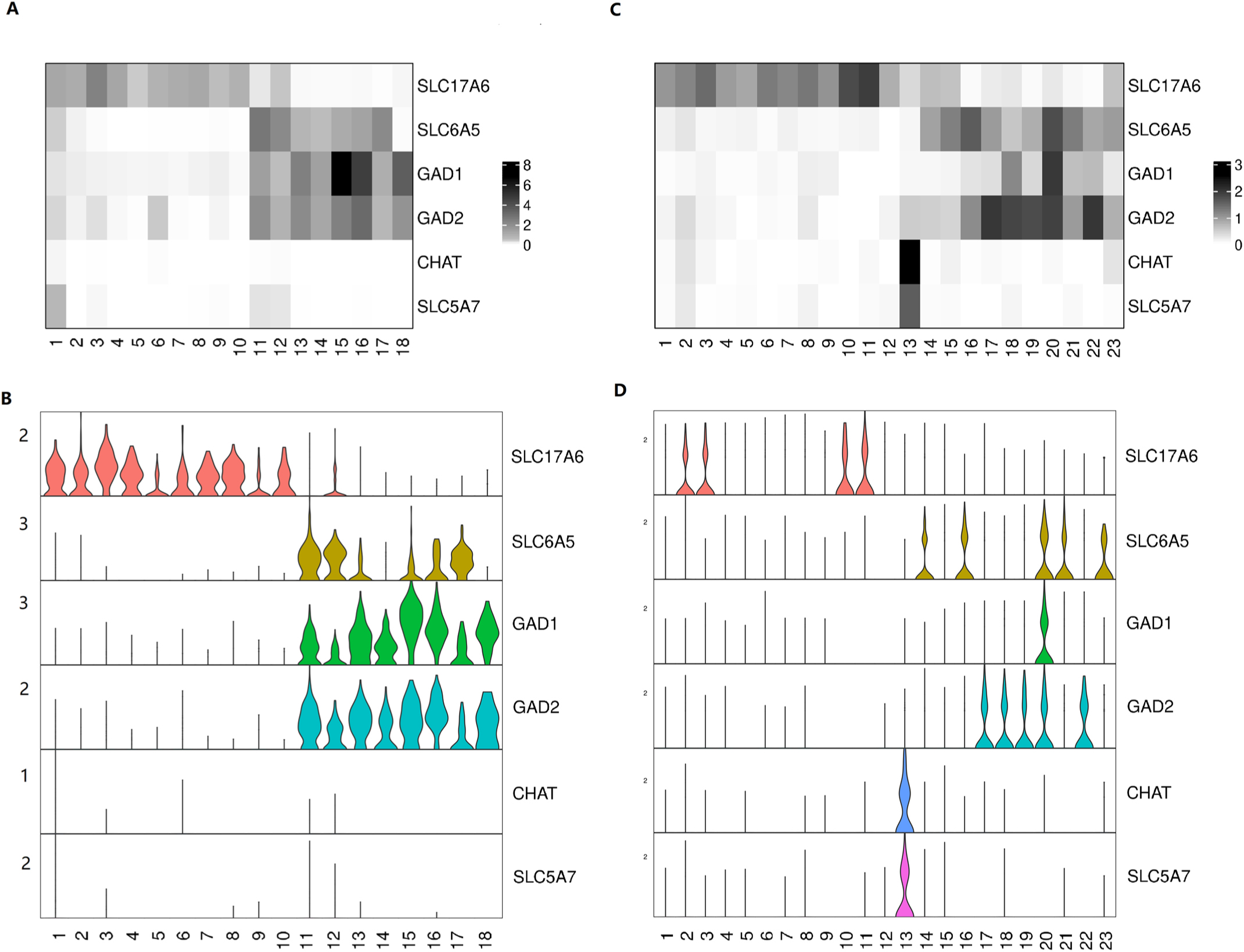
(A-B) Violin plot (A) and heatmap (B) showing the distribution of expression levels of excitatory and inhibitory marker genes across all 18 neuronal clusters in human spinal cord. (C-D) Violin plot (A) and heatmap (B) showing the distribution of expressional levels of excitatory and inhibitory marker genes across all the 23 neuronal clusters in mouse spinal cord.

**Supplementary Figure 3.**
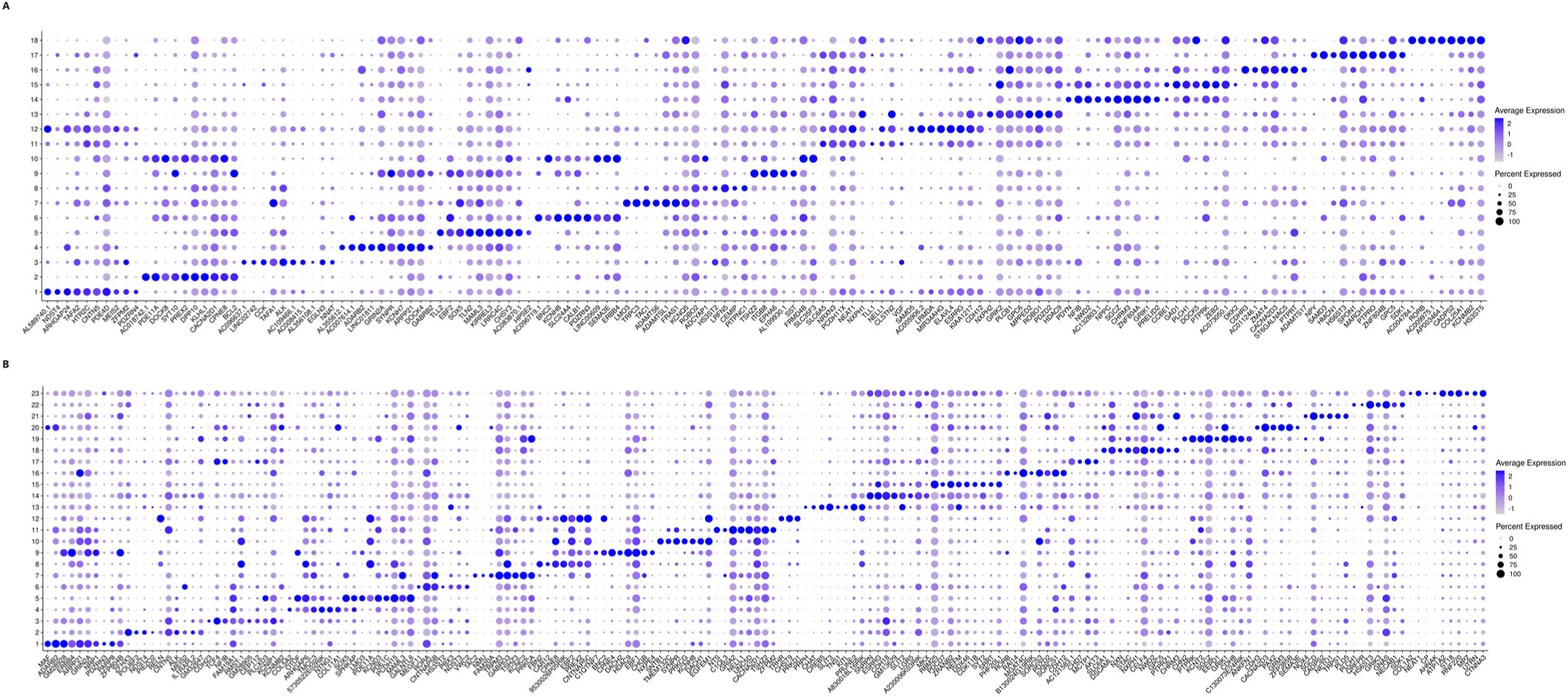
(A-B) Dot plot showing the expression of the top ten most differentially expressed genes across all the neuronal clusters of spinal cord in human (A) and mouse (B).

**Supplementary Figure 4.**
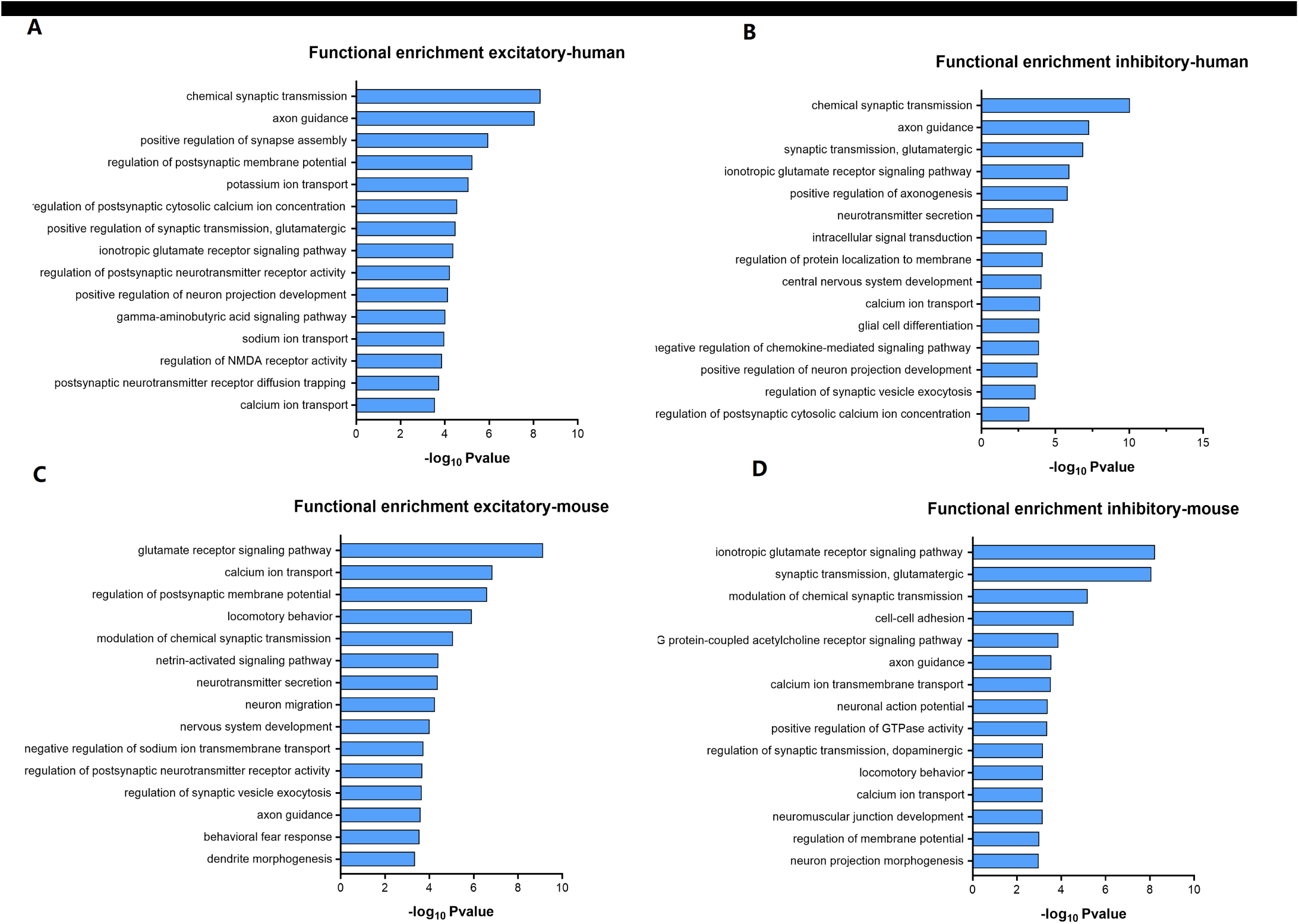
(A-B) Summarized GO terms among the top genes in excitatory (A) and inhibitory (B) clusters of human spinal cord. (C-D) Summarized GO terms among the top genes in excitatory (C) and inhibitory (D) clusters of mouse spinal cord.

**Supplementary Figure 5.**
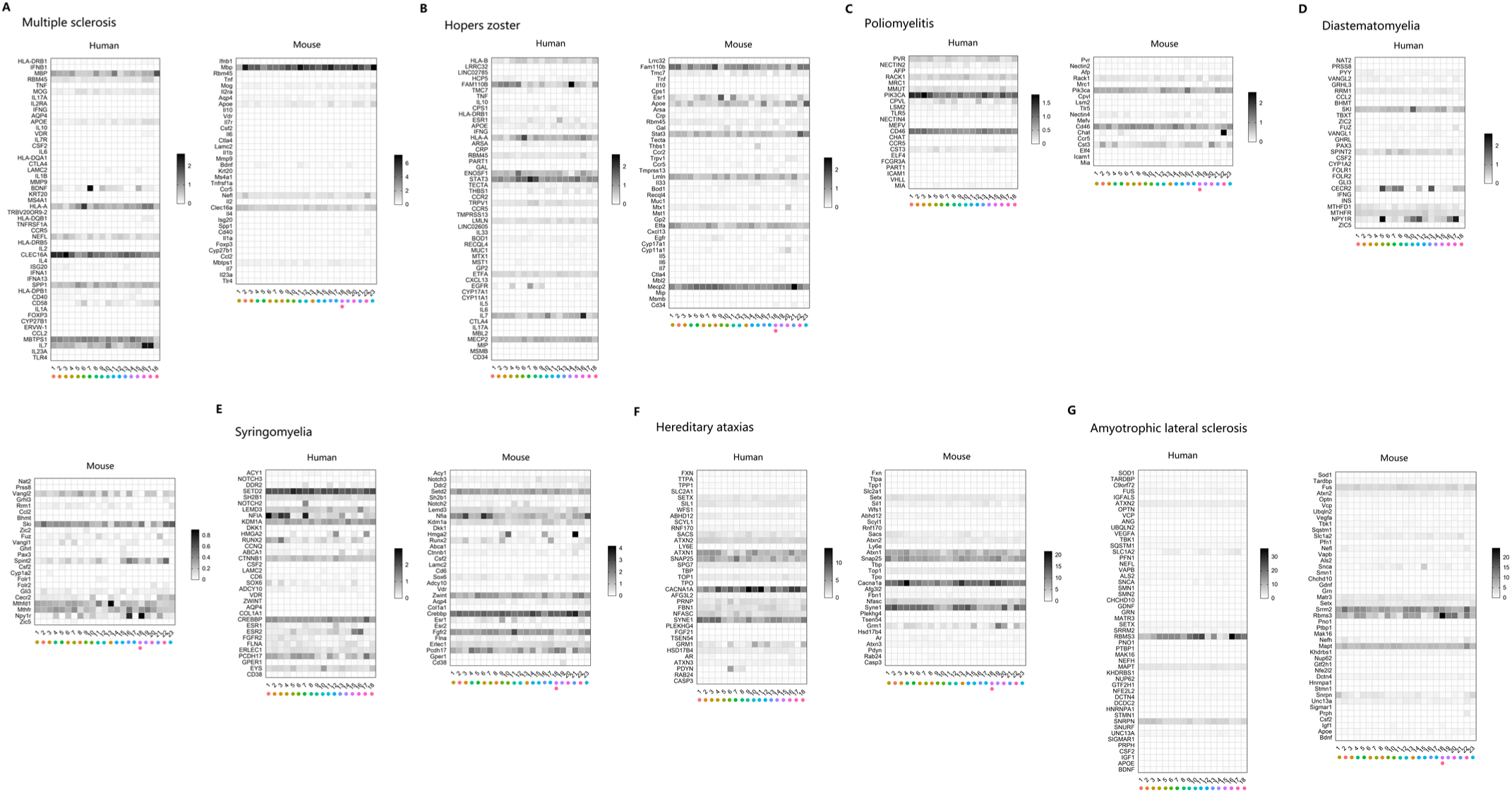
(A-G) The transcriptional profiles of risk genes for Multiple sclerosis (A), herpes zoster (B), poliomyelitis (C), Diastematomyelia (D), Syringomyelia (E), Hereditary ataxias (F), and Amyotrophic lateral sclerosis (G).

**Supplementary Figure 6.**
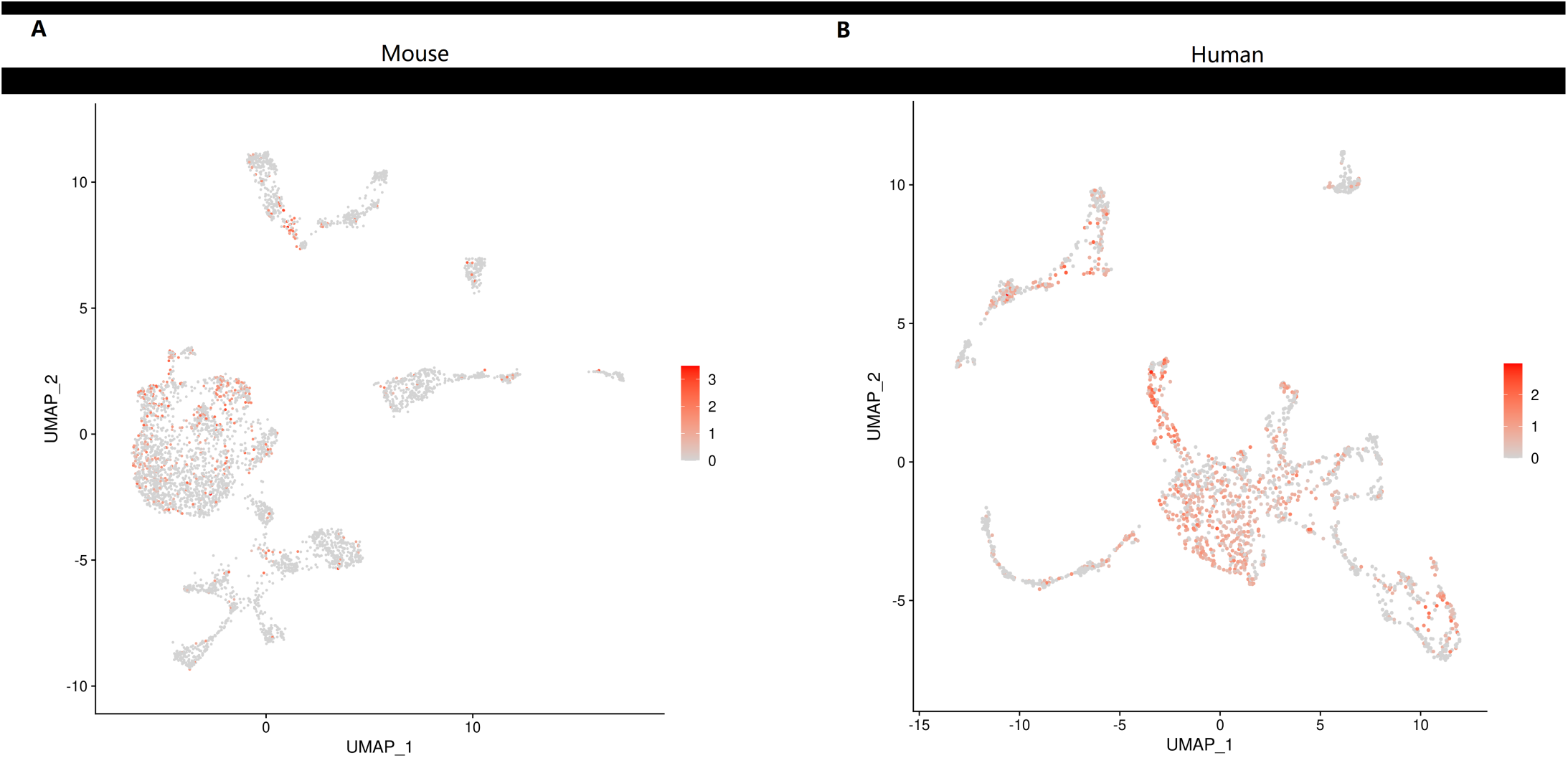
(A-B) UMAP plot showing the distribution of expression of TACR1 across all neuronal types in the spinal cord of mouse (A) and human (B).

## References

Abraira, V.E., Kuehn, E.D., Chirila, A.M., Springel, M.W., Toliver, A.A., Zimmerman, A.L., Orefice, L.L., Boyle, K.A., Bai, L., Song, B.J., et al. (2017). The Cellular and Synaptic Architecture of the Mechanosensory Dorsal Horn. Cell 168, 295–310.e219.

Aldinger, K.A., Thomson, Z., Phelps, I.G., Haldipur, P., Deng, M., Timms, A.E., Hirano, M., Santpere, G., Roco, C., Rosenberg, A.B., et al. (2021). Spatial and cell type transcriptional landscape of human cerebellar development. Nat Neurosci 24, 1163–1175.

Bourane, S., Grossmann, K.S., Britz, O., Dalet, A., Del Barrio, M.G., Stam, F.J., Garcia-Campmany, L., Koch, S., and Goulding, M. (2015). Identification of a spinal circuit for light touch and fine motor control. Cell 160, 503–515.

Delile, J., Rayon, T., Melchionda, M., Edwards, A., Briscoe, J., and Sagner, A. (2019). Single cell transcriptomics reveals spatial and temporal dynamics of gene expression in the developing mouse spinal cord. Development 146.

Grindberg, R.V., Yee-Greenbaum, J.L., McConnell, M.J., Novotny, M., O’Shaughnessy, A.L., Lambert, G.M., Araúzo-Bravo, M.J., Lee, J., Fishman, M., Robbins, G.E., et al. (2013). RNA-sequencing from single nuclei. Proc Natl Acad Sci U S A 110, 19802–19807.

Habib, N., Avraham-Davidi, I., Basu, A., Burks, T., Shekhar, K., Hofree, M., Choudhury, S.R., Aguet, F., Gelfand, E., Ardlie, K., et al. (2017). Massively parallel single-nucleus RNA-seq with DroNc-seq. Nat Methods 14, 955–958.

Ju, G., Hökfelt, T., Brodin, E., Fahrenkrug, J., Fischer, J.A., Frey, P., Elde, R.P., and Brown, J.C. (1987). Primary sensory neurons of the rat showing calcitonin gene-related peptide immunoreactivity and their relation to substance P-, somatostatin-, galanin-, vasoactive intestinal polypeptide-and cholecystokinin-immunoreactive ganglion cells. Cell Tissue Res 247, 417–431.

Kushnarev, M., Pirvulescu, I.P., Candido, K.D., and Knezevic, N.N. (2020). Neuropathic pain: preclinical and early clinical progress with voltage-gated sodium channel blockers. Expert Opin Investig Drugs 29, 259–271.

Lake, B.B., Ai, R., Kaeser, G.E., Salathia, N.S., Yung, Y.C., Liu, R., Wildberg, A., Gao, D., Fung, H.L., Chen, S., et al. (2016). Neuronal subtypes and diversity revealed by single-nucleus RNA sequencing of the human brain. Science 352, 1586–1590.

Li, C.L., Li, K.C., Wu, D., Chen, Y., Luo, H., Zhao, J.R., Wang, S.S., Sun, M.M., Lu, Y.J., Zhong, Y.Q., et al. (2016). Somatosensory neuron types identified by high-coverage single-cell RNA-sequencing and functional heterogeneity. Cell Res 26, 83–102.

Li, L., Rutlin, M., Abraira, V.E., Cassidy, C., Kus, L., Gong, S., Jankowski, M.P., Luo, W., Heintz, N., Koerber, H.R., et al. (2011). The functional organization of cutaneous low-threshold mechanosensory neurons. Cell 147, 1615–1627.

Mishra, S.K., and Hoon, M.A. (2013). The cells and circuitry for itch responses in mice. Science 340, 968–971.

Perez-Cruz, F. (2008). Kullback-Leibler divergence estimation of continuous distributions. Paper presented at: 2008 IEEE International Symposium on Information Theory.

Rosenberg, A.B., Roco, C.M., Muscat, R.A., Kuchina, A., Sample, P., Yao, Z., Graybuck, L.T., Peeler, D.J., Mukherjee, S., Chen, W., et al. (2018). Single-cell profiling of the developing mouse brain and spinal cord with split-pool barcoding. Science 360, 176–182.

Sathyamurthy, A., Johnson, K.R., Matson, K.J.E., Dobrott, C.I., Li, L., Ryba, A.R., Bergman, T.B., Kelly, M.C., Kelley, M.W., and Levine, A.J. (2018). Massively Parallel Single Nucleus Transcriptional Profiling Defines Spinal Cord Neurons and Their Activity during Behavior. Cell Rep 22, 2216–2225.

Stuart, T., Butler, A., Hoffman, P., Hafemeister, C., Papalexi, E., Mauck, W.M., 3rd, Hao, Y., Stoeckius, M., Smibert, P., and Satija, R. (2019). Comprehensive Integration of Single-Cell Data. Cell 177, 1888–1902.e1821.

Tibbs, G.R., Posson, D.J., and Goldstein, P.A. (2016). Voltage-Gated Ion Channels in the PNS: Novel Therapies for Neuropathic Pain? Trends Pharmacol Sci 37, 522–542.

Todd, A.J. (2010). Neuronal circuitry for pain processing in the dorsal horn. Nat Rev Neurosci 11, 823–836.

Todd, A.J. (2017). Identifying functional populations among the interneurons in laminae I-III of the spinal dorsal horn. Mol Pain 13, 1744806917693003.

Yekkirala, A.S., Roberson, D.P., Bean, B.P., and Woolf, C.J. (2017). Breaking barriers to novel analgesic drug development. Nat Rev Drug Discov 16, 545–564.

